## Supplementary figures and images for "Nifuroxazide Suppresses PD-L1 Expression and Enhances the Efficacy of Radiotherapy in Hepatocellular Carcinoma"

### Figure 1-figure supplement 1

Supplementary Fig. 1

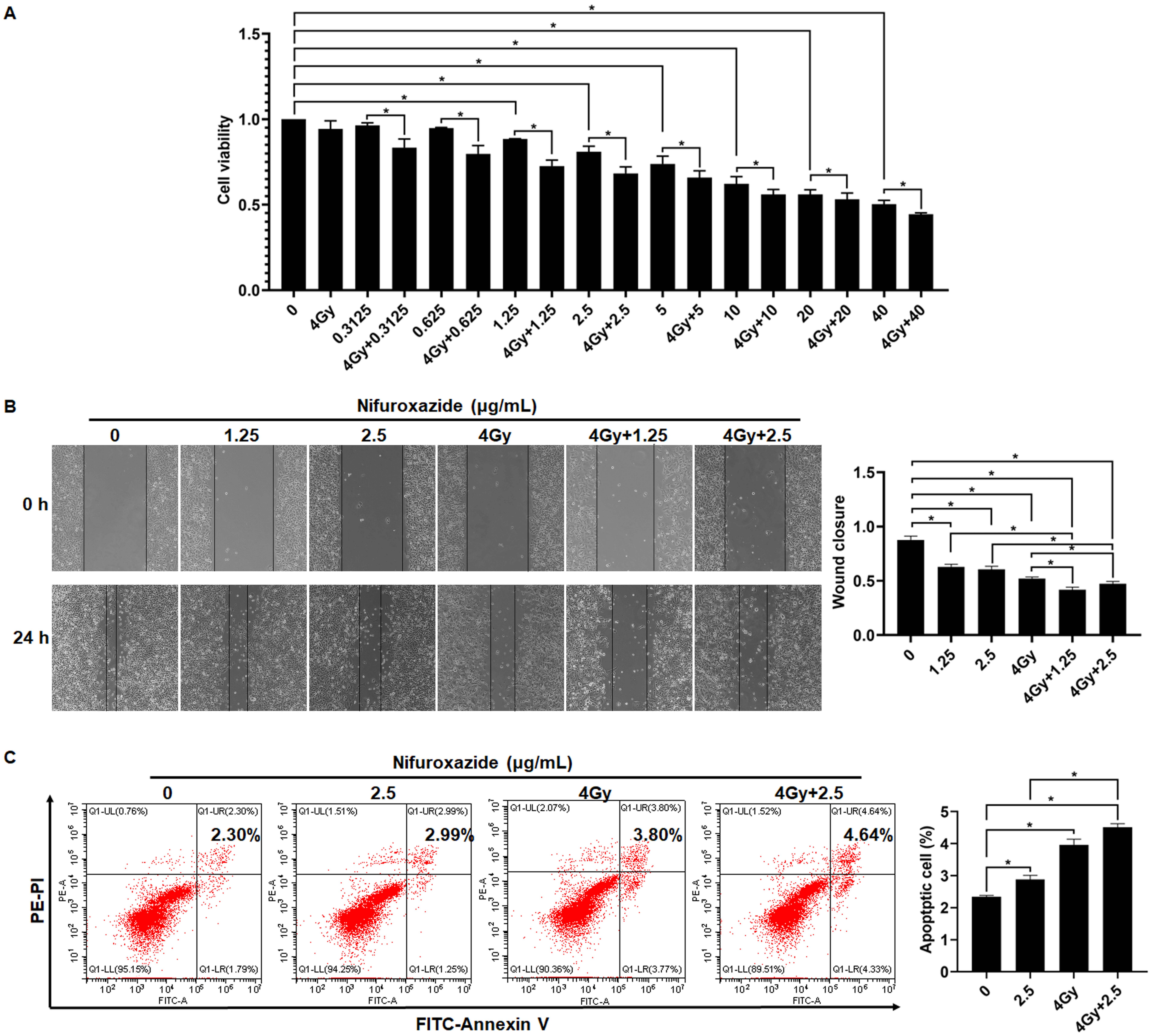

### Figure 5-figure supplement 2

Supplementary Fig. 2

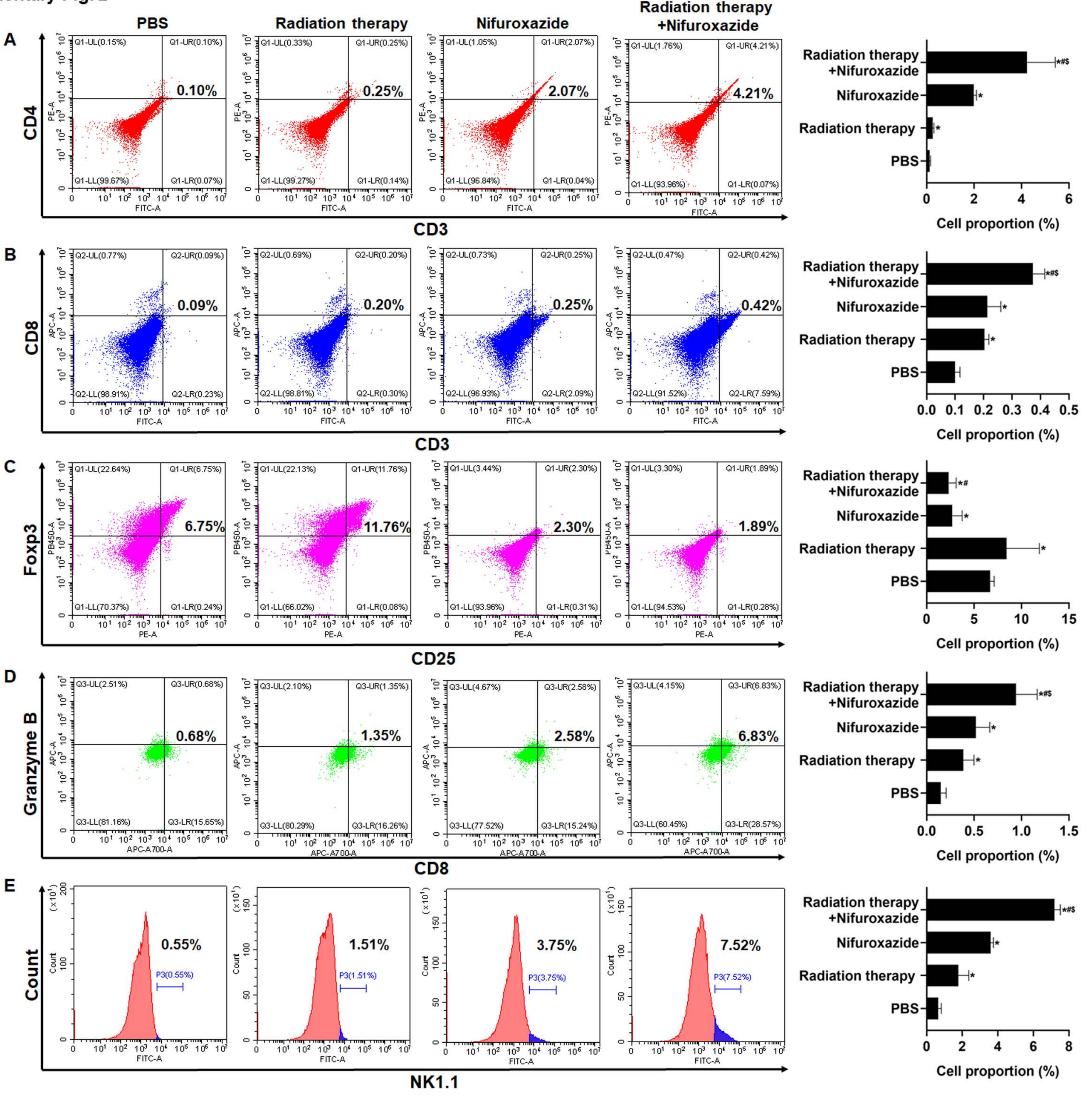

### Figure 7-figure supplement 3

Supplementary Fig. 3

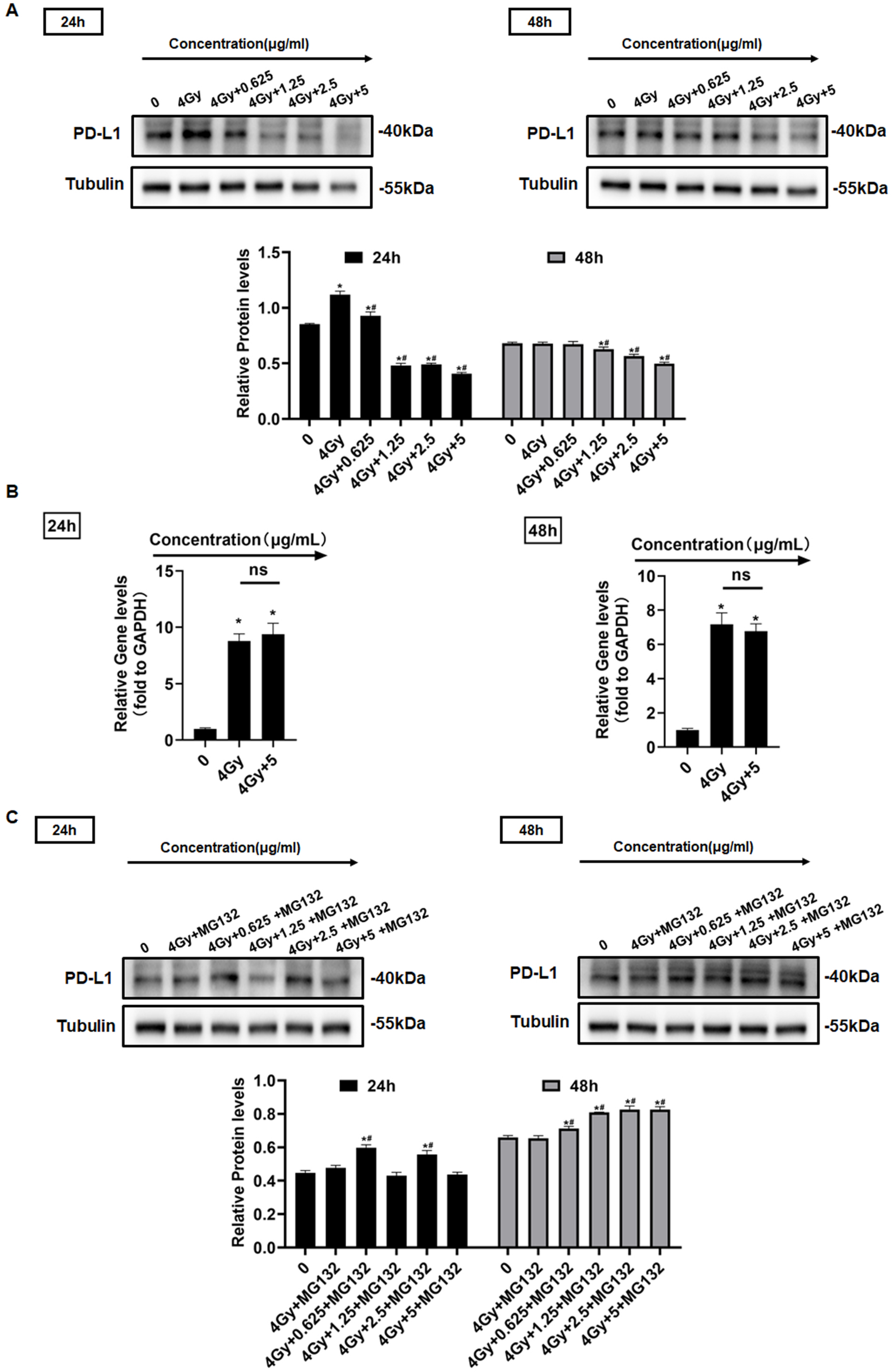
